# Tracking *Microcystis* viruses and infection dynamics across distinct phases of a *Microcystis*-dominated bloom

**DOI:** 10.1101/2024.05.24.595742

**Authors:** A.J Wing, Bridget Hegarty, Eric Bastien, Vincent Denef, Jacob Evans, Gregory Dick, Melissa Duhaime

## Abstract

Given the impact of viruses on microbial community composition and function, viruses have the potential to play a significant role in the fate of freshwater cyanobacterial harmful algal blooms (cHABs). Yet the role of viruses in cHABs remains poorly understood. We sought to address this knowledge gap with a metagenomic analysis of viruses of bloom-forming *Microcystis aeruginosa* across cHAB phases in the western basin of Lake Erie. Size-fractionation of the water allowed us to identify significant fraction-specific trends in viral diversity, which corresponded with *Microcystis* genetic diversity. Using a new machine-learning model, we predicted infections between viral and microbial host populations. We predicted hundreds of viral populations with infection histories including *Microcystis* and non-*Microcystis* hosts, suggesting extensive interconnectivity and the potential for virus-mediated cross-species exchange of genetic material within cHABs communities. Infection predictions revealed a broad host range for Lake Erie *Microcystis* viruses, challenging previous notions of “narrow” host-virus interactions in cHABs. Abundant viral genes belonging to predicted *Microcystis* viruses revealed their potential role in key metabolic pathways and adaptation to environmental changes. We observed significant turnover of predicted *Microcystis* virus populations across time. Viral diversity was highest in the viral fraction and lowest in the colony-associated fraction, suggesting that *Microcystis* colony formation and growth during cHABs leads to bottlenecks in viral diversity. These findings advance our understanding of uncultivated *Microcystis* virus diversity, their potential effects on host metabolism, potential influence on species interactions, and potential coevolutionary processes between microbial hosts and their viral predators within *Microcystis*-dominated cHABs.

**Importance:** Understanding interactions between viruses, their hosts, and environmental parameters may be key to identifying the mechanisms underlying the persistence and demise of cyanobacterial harmful algal blooms (cHABs). In this study we describe the viral diversity and host ranges of viruses predicted to infect *Microcystis*, describing the distribution of these properties across time, space, and different bloom-associated size fractions. Additionally, the study highlights abundant genes belonging to predicted *Microcystis* viruses and their potential roles in key metabolic pathways and adaptation to environmental changes. The observed turnover of *Microcystis* virus populations, with the highest diversity in viral fractions and the lowest in colony-associated fractions, suggests that *Microcystis* colony formation during blooms plays an important role in shaping viral diversity and community turnover. These findings contribute to a better understanding of the interplay between viruses, *Microcystis*, and their accompanying bacterial communities, shedding light on mechanisms driving bloom dynamics, species interactions, and coevolutionary processes.

## Introduction

*Microcystis aeruginosa* is a cyanobacterium that can form toxic blooms in freshwater and estuarine systems worldwide.^67,94^ Microcystins, the most prolific in a suite of toxins produced by *M. aeruginosa,*^64^ have toxic effects on humans and diminish drinking water quality and overall aquatic ecosystem health.^25,61,80^ The frequency and intensity of *M. aeruginosa* blooms are increasing,^20,62^ largely due to climate change and eutrophication of aquatic habitats.^54,60,61,62,84^ While the role of such abiotic controls on *Microcystis* bloom progression has been the focus of study for years, elucidating the role of biotic controls, such as predator-prey relationships, has been less explored, owing partly to the challenges of studying microbial interactions in complex community contexts.

Viruses profoundly influence microbial communities by infecting and lysing microbial host populations,^17,40,82,87^ reprogramming host metabolisms,^7,15,24,27,70,98^ and facilitating gene transfer.^52,79^ Evidence suggests that *Microcystis* in the annual Lake Erie cHABs of the lake’s western basin is susceptible to viral infection and subsequent lysis.^33,53,80^ Release of microcystins from cells due to viral lysis contributed to the Toledo drinking water crisis in 2014.^80^ Yet, much of what is known of *Microcystis* viruses is through tracking the abundance and distribution of viral marker genes of a single *Microcystis* virus isolate, Ma-LMM01, and its close relatives.^51,53,74,80,95,96^ In contrast, a community genomic approach captures the complex system of microbial populations interacting with one another, their viral predators, and their environment.

In this study, we sought to address (1) How do the diversity and distribution of viruses infecting *Microcystis* vary across time, space, and different size fractions during a cHAB, and (2) What are the possible implications of *Microcystis* virus infection in terms of host metabolisms and gene flow with other host taxa? With a combination of metagenomic analyses of viral and cellular communities, we set out to detail the viral diversity and potential host range of viruses predicted to infect *Microcystis*, the distribution of these viruses across time, space, and different size fractions, and explored their metabolic potential.

## Results and Discussion

### Tracking *Microcystis* during the 2014 Lake Erie cHAB

We analyzed metagenomic data obtained from cyanobacterial harmful algal blooms (cHABs) in the western region of Lake Erie (Fig. 1A) that persisted from July to October in 2014.^4,13,78^ Concentrations of particulate phycocyanin (used as a proxy for cyanobacteria) and microcystin (indicative of bloom toxicity) revealed a toxic *Microcystis* bloom at all three sampling locations (WLE12, WLE2, WLE4) in early August (Fig. 1A-B). Nearshore stations WLE12 and WLE2 consisted of *Microcystis* genotypes with complete mcy operons, the gene cassette responsible for microcystin production, during early August, whereas the *Microcystis* population at offshore station WLE4 contained a mixture of complete and partial mcy operons.^92^ The cHAB in late September occurred primarily at the nearshore stations with lower microcystin concentrations than August 4 (Fig. 1B); the *Microcystis* genotypes in this second bloom lacked complete mcy operons or lacked mcy genes altogether.^92^

**Figure 1.**
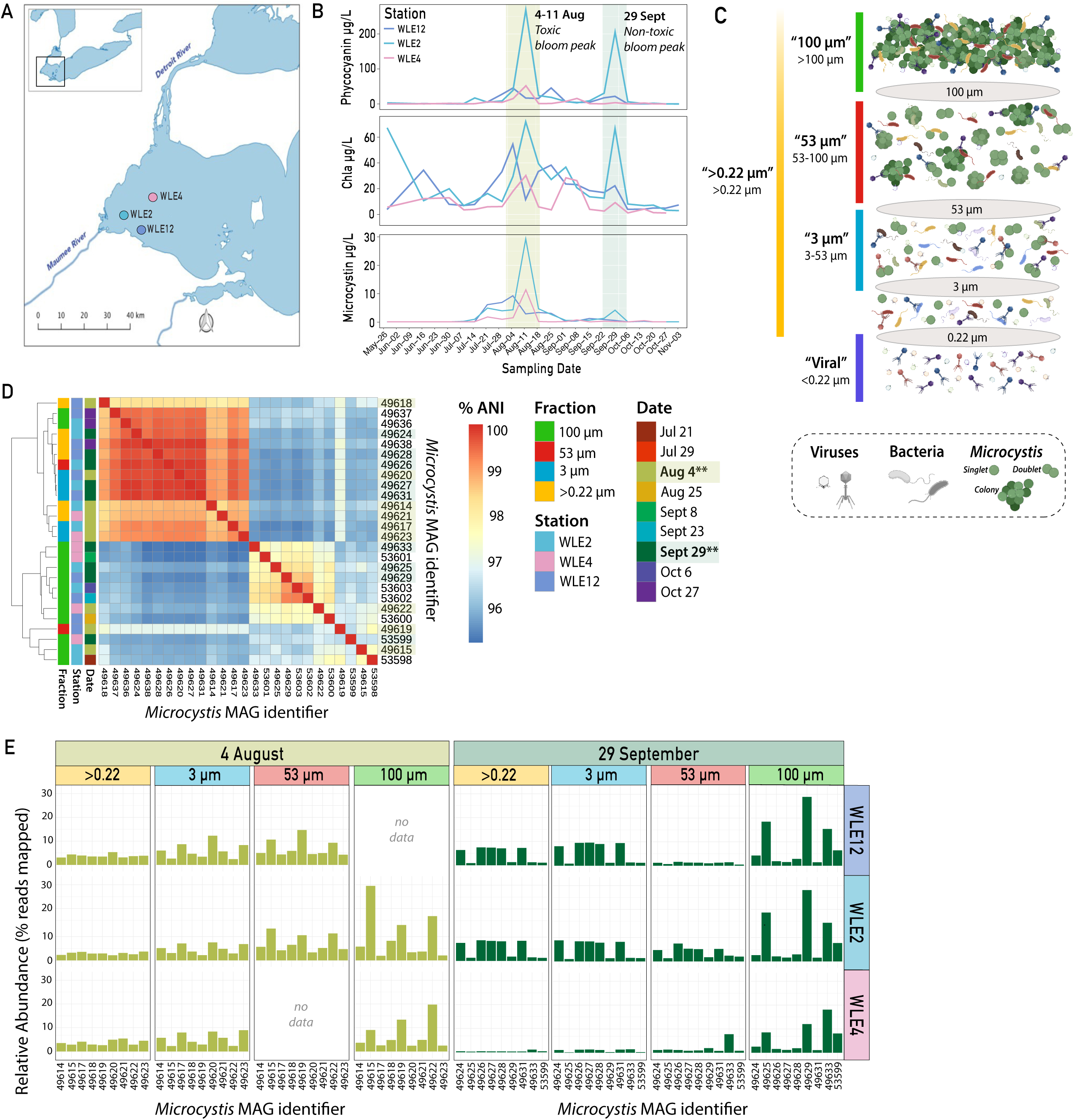
Sampling overview and *Microcystis* MAG diversity and dynamics. (A) Map of sampling sites in the western basin of Lake Erie. Three sites were sampled bi-monthly in June and weekly from July to October of 2014. (B) Phycocyanin, chlorophyll-a and microcystin measurements for the 2014 bloom. (C) Graphical summary of sampling filter size fractionation depicting how samples were collected. Viruses, non-*Microcystis* bacteria, and *Microcystis* are depicted by virions, multicolored flagellated rods, and green cocci (as singlets, doublets, and colonies), respectively. (D) Heatmap of average nucleotide identity (% ANI) between 26 *Microcystis* MAGs reconstructed from samples from July through August across three stations and four sample fractions. Bottom row and right column list the MAG identifiers. Sample fraction, station and date are identified by color of left panel. Dendrogram depicts the clustering of MAGs based on %ANI. (E) Relative abundances of *Microcystis* MAGs based on fraction of reads from each sample that mapped to each MAG in a competitive read mapping to all assembled MAGs from that sample. No data exist for 4 Aug 100 µm at WLE12 and 53 µm at WLE4.

To investigate the viruses and virus-host interactions associated with these blooms, we analyzed metagenomic data from each station at the two bloom peaks and across five size fractions chosen to target *Microcystis* in both free-living and colony forms and their viruses (Fig. 1C). From these data, we reconstructed 41 bacterial metagenome assembled genomes (‘MAGs’), 17 of which were taxonomically classified as *Microcystis* (Fig. 1D-E; SI Table 1) and 27,086 viral contigs >3 kb. Relative abundances of *Microcystis* ranged from 0-29%, based on the proportion of the total reads in each sample that mapped to the *Microcystis* MAGs in the cellular fraction metagenomes (0.22-100 µm; Fig. 1E). Highest relative abundances of *Microcystis* MAGs were observed in the 100 µm fraction where large *Microcystis* colonies that typify blooms are expected. The greatest variability in abundances of individual *Microcystis* MAGs was also observed in this fraction (Fig. 1E), which we suspect is due to the sporadic rise and fall of different colony-forming *Microcystis* populations captured in the 100 µm fraction. Viral contigs identified across all size fractions were clustered into 15,461 viral operational taxonomic units (vOTUs), which approximate viral species, based on broadly accepted thresholds (95% ANI across 85% of the contig length).^72^

### Microbial virus-host interaction networks at toxic and nontoxic bloom peaks

#### Hundreds of viral OTUs predicted to infect *Microcystis* at bloom peaks

We sought to identify likely infection linkages between uncultivated viruses and *Microcystis* population genomes (MAGs) reconstructed from the two 2014 Lake Erie cHAB bloom peaks. We used Virus-Host Interaction Predictor (VHIP),^3^ a machine learning-based infection prediction tool that predicts putative virus-host pairs. The approach leverages genome-encoded signals of coevolution (e.g., nucleotide frequencies, %G+C patterns, shared nucleotide and protein sequences, etc.) in a model trained and tested on 8,849 lab-verified infection/no-infection data. We expanded the VHIP analysis to also include non-*Microcystis* MAGs that were predicted to be infected by identified *Microcystis* viruses.

At the August 4 toxic bloom peak, a total of 2,026 virus-host pairs were predicted between 454 vOTUs and 17 bacterial MAGs (9 of which were *Microcystis* MAGs; SI Table 1) (Fig. 2A). On the September 29 non-toxic bloom peak, 1,995 virus-host pairs were predicted between 339 vOTUs and 24 bacterial MAGs (8 *Microcystis* MAGs; SI Table 1) (Fig. 2B). Moreover, VHIP predicted a higher number of non-infections compared to infections, where most predictions were heavily skewed towards either strong signals of non-infection or infection (SI Fig. S1). Notably, the majority of viruses were predicted to strongly infect at least one host, while others were predicted to be incapable of infecting any host (SI Fig. S2). The majority of viruses predicted to infect *Microcystis* were present at low abundances in the bloom peak samples, particularly on August 4. Abundant vOTUs (vOTUs that recruited >0.1% of the total reads mapped to vOTUs) represented 6.6% and 13.9% of the total *Microcystis* vOTUs on 4 Aug and 29 Sept, respectively. This trend of majority low abundance and few high abundance populations is common in microbial virus assemblages, which tend to have long tailed rank-abundance curves like their bacterial and archaeal hosts.^10,14,49^ The top most numerically abundant *Microcystis* vOTUs that emerged on the bloom peaks (labeled in Fig. 2A-B) are discussed in the context of assemblage evenness and turnover in later sections.

**Figure 2.**
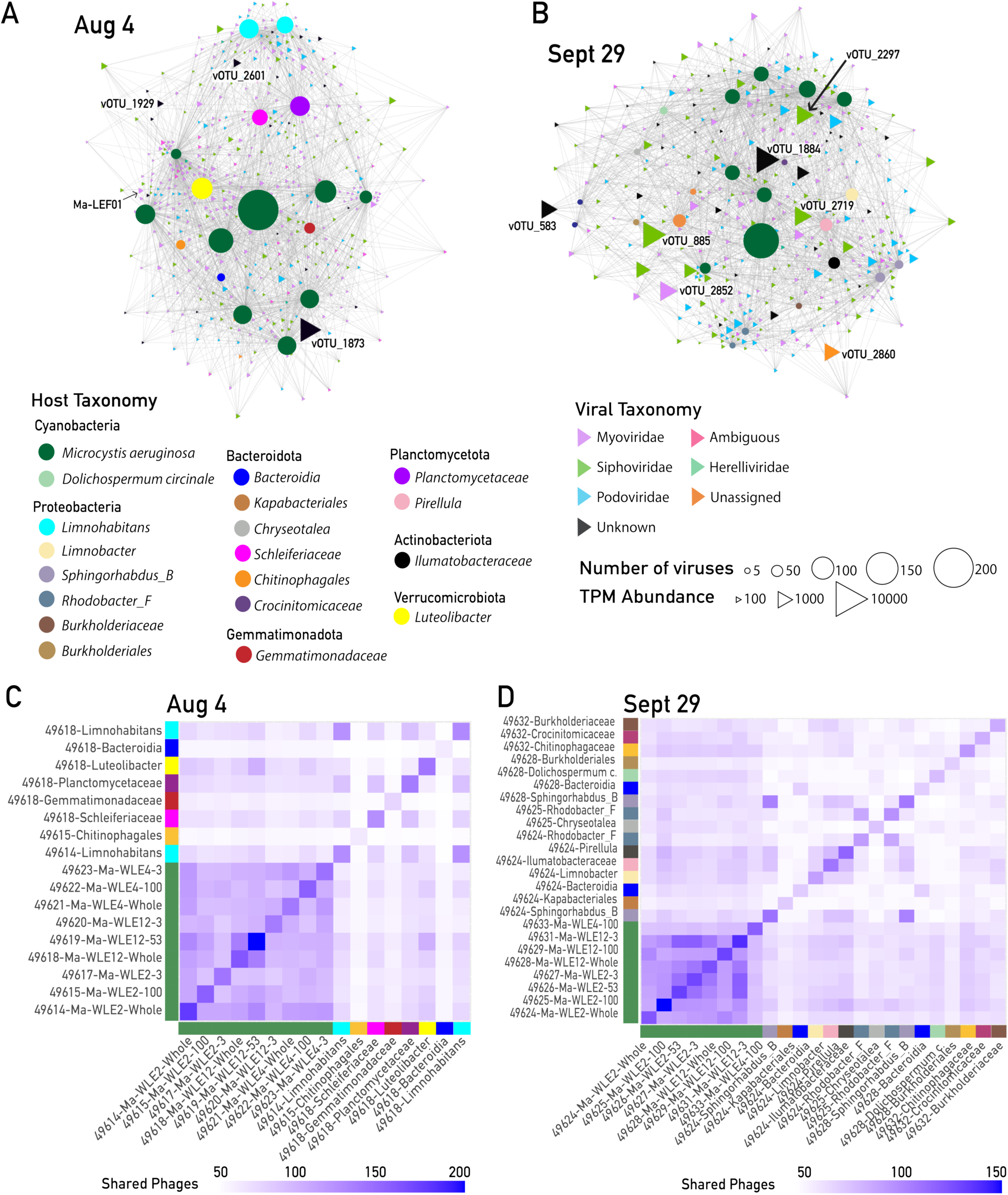
Networks of predicted infections between *Microcystis* viruses (>10 kb viral contigs) and bacterial host MAGs identified on the 4 Aug and 29 Sept bloom metagenomes. (A) Predicted infection network of 4 August toxic bloom peak. (B) Predicted infection network of 29 September non-toxic bloom peak. Circle nodes are host MAGs; circle size represents the number of viruses predicted to infect a given host. Triangle nodes are viruses. Node size represents TPM abundance. Node colors represent assigned taxonomy; for viruses, “Unknown” indicates the vOTU has no hit in the reference database, “Unassigned” indicates the vOTU has a hit to something unassigned in the reference database. Only predictions with >93% infection probability and viruses predicted to infect at least *Microcystis* are shown. The most abundant vOTUs are labeled in each network (described in SI Table 2 and Fig. 4) (C) Heat map of hosts with shared virus predictions on 4 August. (D) Heat map of hosts with shared virus predictions on 4 August. Axis colors represent host taxonomy (as in panels A-B) and heat map cell color represents the amount of shared phages between any two given hosts.

While we have no available methods to determine the true number of uncultivated viruses infecting a given uncultivated host, these numbers of predicted *Microcystis* viruses are higher than previously reported from metagenomic studies that used other approaches to identify putative *Microcystis*-infecting viruses.^53,56,66^ Given the 87.8% accuracy of VHIP in predicting infections,^3^ we are confident that a majority of the co-evolutionary associations identified in these data reflect true virus-host interactions. CRISPR spacer-only infection networks predicted only 16 infections total across two bloom peaks, far fewer than the VHIP network (SI Fig. S3). These CRISPR spacer-only predictions accounted for only 0.004% of all VHIP predictions. The elevated number of predicted *Microcystis* vOTUS may be partially attributed to the incomplete nature of metagenome-reconstructed population genomes. Genomic “shrapnel,” which are genome fragments that could not be linked with their corresponding parts due to the challenges of binning viral genomes,^35,73^ likely elevate the number of predicted infections (e.g., one virus could be counted as many). The mean viral bin length in this study was 32.7 kb, compared to the mean NCBI bacteriophage genome length of 68.4 kb, suggesting that incomplete viral genome assembly and binning may contribute at least a 2-fold overestimation of phage-host “linkages” in the network. While the numbers of predicted viruses are not likely to reflect the true number of viruses in a given sample owing to caveats described below, the overall data structures are informative. These linkages provide novel perspectives regarding the structure of the predator-prey networks (e.g., narrow versus broad host ranges), the potential for gene flow between host and virus populations (e.g., *“which host populations are or have been evolutionarily connected via viral infection?”*), and insights into how viral diversity fluctuates through time and space over the course of a bloom.

#### Considerations on linking uncultivated viral and host populations using genomic signals of coevolution

Our method for predicting bacterial hosts of putative *Microcystis* viruses overcomes the limitations of marker gene analyses, which have historically been restricted to tracking and linking a single viral population,^39,51,53,66,83,94^ and CRISPR-based approaches,^55,56^ which are limited by the absence of detectable CRISPR systems in many bacterial genomes.^9^ Additionally, our method does not rely on coassociations between viral and *Microcystis* marker genes,^66^ which may oversimplify ecological complexities by assuming binary interactions and linear correlations, when in reality these ecological relationships are varied and rarely linear.^12,23^

While interpreting these VHIP-generated networks, it is helpful to consider them as a superposition of all theoretically possible infection networks, rather than as a snapshot of active infections captured at the time and place of sampling. Because VHIP relies on signals of coevolution detected between viral and putative host genomes and because different coevolutionary signals establish and degrade at different rates, there will be remnant signals linking populations that may not be able to carry out infections—for instance, because their populations may never coexist in time and space. Nonetheless, the model predicts lab-verified infection pairs of species-level virus-host linkages with 87.8% accuracy.^3^ We applied VHIP to gain a better understanding of the diversity, turnover, and metabolic impacts of *Microcystis* viruses at the bloom peaks.

#### Most *Microcystis* vOTUs host ranges are within-genus, some span phyla

To evaluate *Microcystis* vOTU host range and potential for virus-mediated cross-host gene transfer, we identified the predicted *Microcystis* vOTUs that were also predicted to infect non-*Microcystis* MAGs. Whereas the vast majority of shared viruses were between *Microcystis* populations (Fig. 2C-D), eight and 16 non-*Microcystis* hosts on Aug 4 and Sept 29, respectively, were predicted to share viruses with *Microcystis* (Fig. 2). Most notably, on August 4, five prominent host nodes representing four non-Cyanobacteria phyla had a relatively high number of *Microcystis* viruses predicted to infect them (Fig. 2A). While *Microcystis* viruses have been shown to infect multiple cyanobacterial genera in lab studies,^86^ tests across higher phylogenetic levels have not been reported. However, viral isolates of other host taxa *are* known to infect across multiple phyla;^50^ we suspect the dearth of cross-phyla reports is not only due to evolutionary constraints that underlie host range, but also because of how extremely rare it is for cross-phyla infections to be tested in the laboratory.^3^

A previous study using qPCR to track metagenome-identified viral groups showed how narrow host range *Microcystis* viruses were observed at markedly lower abundances than broad host range viruses, which tended to be more dominant.^56^ We tested whether similar trends were observed in the Lake Erie bloom. The predicted host range depended on whether only *Microcystis* hosts were considered (as in the Morimoto et al., 2023 study) or all hosts were considered (Fig. 3A). Considering only *Microcystis* hosts, most vOTUs were predicted to infect only one host at the bloom peak and 60% of the viruses were predicted to infect only one or two other *Microcystis* MAGs (Fig. 3A); these would be classified as “narrow” host range viruses by Morimoto et al., 2023. However, considering all hosts, three hosts were predicted per vOTU, with 76% of vOTUs belonging to Morimoto’s “broad” host range category (Fig. 3A). In contrast to Morimoto et al., 2023, we found no or only extremely weak correlations between vOTU host range breadth (i.e., number of hosts) and abundance (Pearson R^2^ <0.05 for all tests; SI Table 3).

**Figure 3.**
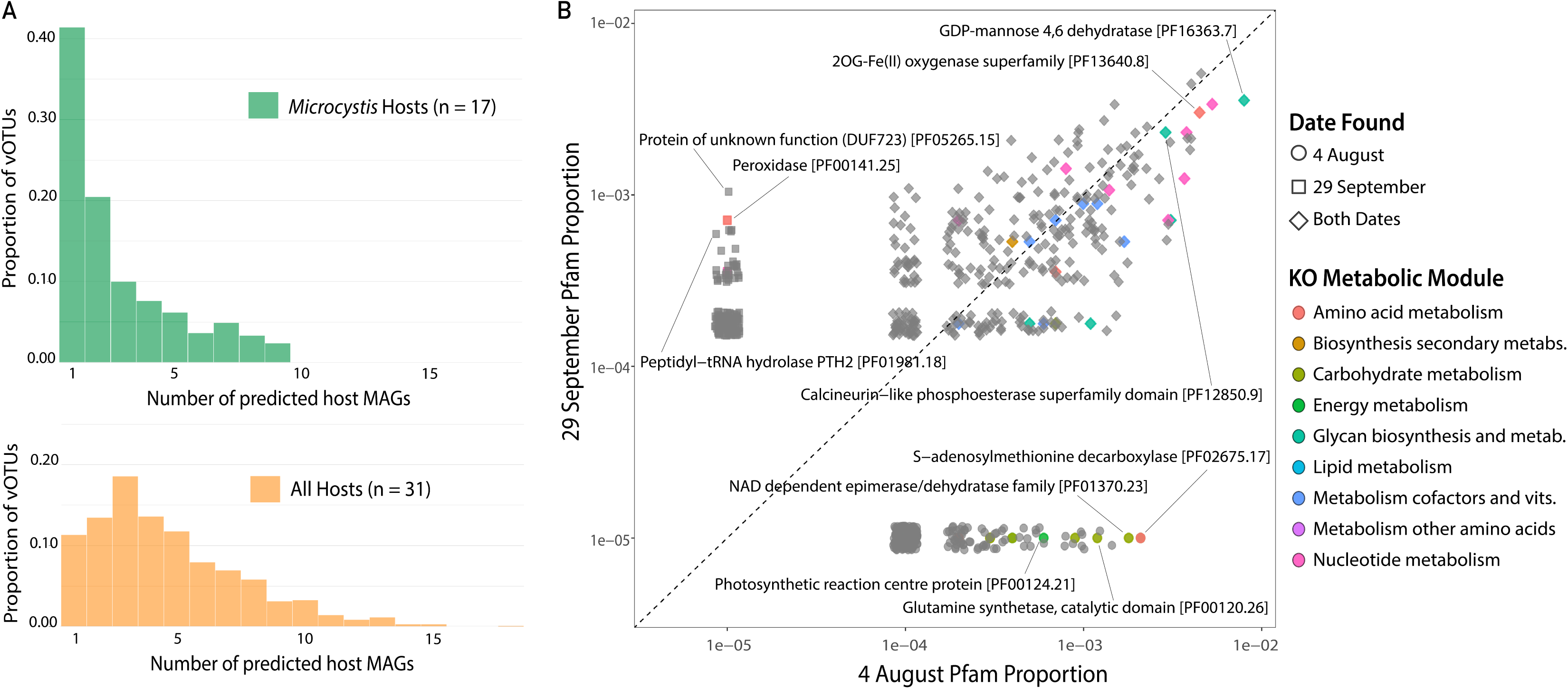
Proportion of vOTUs infecting bloom peak hosts and proportion of abundant viral genes at bloom peaks. (A) Comparison of host ranges. (B) Proportion of viral genes encoded by predicted *Microcystis* vOTUs at 4 August bloom peak (x-axis) and 29 September bloom peak (y-axis). Shape indicates which samples viral genes were identified in. Point color reflects the assigned KEGG Ontology (KO) metabolic module. Points labels are derived from assigned protein family (Pfam) annotations for select functions of interest.

It is important to consider that the terms “broad” and “narrow” are operational and often defined relative to a given study. In the Morimoto study, despite delineating narrow and broad host range groups, all host diversity was constrained to a single host genus. Here, we report *Microcystis* virus-host pairs with genomic evidence that suggests past infections that span phyla. While rampant recombination can spread these signals across the reticulate phylogeny of viruses, there is also culture-based evidence to support that these signals could reflect true cross-phyla infections and community genomic evidence to support cross-domain infections.^28,50^ Future work should focus on experimentally validating these putative cross-phyla and cross-domain infections of *Microcystis* viruses, assessing the mechanisms and ecological and evolutionary implications of such interactions within *Microcystis*-dominated cHABs.

#### Metabolic genes encoded by predicted *Microcystis* vOTUs can be bloom peak-specific

The coevolutionary signals that underlie the predicted virus-host interaction networks represent paths of potential virus-mediated gene flow between bacterial populations. Viruses are well known to encode genes involved in myriad metabolic processes, e.g., photosynthesis, nitrogen metabolism,^73^ and sulfur metabolism.^2,37^ When viruses facilitate the cross-taxa transfer of genes central to bacterial metabolism, there are potential consequences for the evolutionary trajectory of proteins with important biogeochemical functions.^48^ Further, during infection, virus-encoded auxiliary metabolic genes (AMGs) rewire host metabolisms in ways that influence the flow of matter and energy during infection.^24,98^ To evaluate this potential, we identified the metabolic genes carried by the viruses predicted to interact with *Microcystis*.

Most identified *Microcystis* vOTU genes encode proteins with unknown metabolic functions (Fig. 3B; SI Table 4). Of the AMGs that could be annotated, some were shared at the bloom peaks and some were unique to the peaks on either August 4th or September 29th (Fig 3B). Virus-encoded AMGs shared at the bloom peaks included the biosynthesis of complex carbohydrates important for cell wall biosynthesis and colony formation, cell-to-cell communication, and biofilm formation (GDP-Mannose 4,6 dehydratase),^34^ the production of phycobiliprotein light-harvesting pigments and secondary metabolites (2OG-Fe(II) oxygenase superfamily)^22,32^ and the regulation of phosphate metabolism (2OG-Fe(II) oxygenase superfamily).^57,85^ Virus-encoded functions enriched on August 4th included those central to photosynthesis (photosynthetic reaction center proteins), nitrogen, amino acids, and energy metabolism (Glutamine synthetase; NAD-dependent epimerase/dehydratase),^6^ and the cellular responses to nutrient, light and temperature fluctuations (S-adenosylmethionine decarboxylase).^31,97^ Virus-encoded metabolic functions enriched on September 29th included the viral takeover of host protein synthesis (peptidyl-tRNA hydrolase PTH2 enzymes)^18^ and cellular responses to oxidative stress, which can arise from exposure to high light intensity and reactive oxygen species generated during photosynthesis (peroxidase enzymes).^99^ High abundance of viral genes encoding peroxidase gene *ahpC* on 29 September suggested a potential role of viral AMGs in supplementing *Microcystis* ROS defense and may help explain elevated expression of this gene in these samples.^77^

Overall, the functional gene analysis of the *Microcystis* vOTUs indicated that there are bloom phase-dependent consequences for how viruses may be supplementing or rewiring host metabolisms. Given that these AMGs encode essential functions such as photosynthesis, nutrient metabolism, and oxidative stress response, these results suggest that viruses may be relieving metabolic and physiological bottlenecks, which likely vary across bloom stages. These findings motivate future work to better resolve the degree to which viruses shape the genotypic and phenotypic changes that manifest in the cyanobacterial populations that dominate Lake Erie’s cHABs.

### Turnover of predicted *Microcystis* vOTUs depends on colony formation

#### Diversity of predicted *Microcystis* vOTUs is highest in the viral fraction, lowest in the colony-associated fraction

vOTU abundances estimated based on sequence read recruitment were used to evaluate trends in diversity within the subset of vOTUs predicted to infect *Microcystis*. Within this *Microcystis* vOTU assemblage, evenness was the highest in the virus fraction and decreased with each consecutively larger pore size filter fraction (Fig. 4A). This trend is consistent with observed host strain diversity; the *Microcystis* MAGs were less evenly distributed in the larger colony-associated fraction (100 µm) than the smaller size fractions (>0.22 and 3 µm) (Fig. 1E). We considered 53 and 100 µm to be ‘colony-associated’ fractions, while those ‘not colony-associated’ were viral, 3 µm and >0.22 µm, the latter of which has been shown in other lake systems to be numerically dominated by free-living cells.^75^ The genomic variation observed in the *Microcystis* MAGs from Lake Erie showed a distinct partitioning by size fraction. Such within-species genotypic differences that associate with different filter fractions have been observed in ocean taxa as well, such as *Vibrio splendidus*.^26^ Notably, a significant clustering of genotypes was evident in the 100 µm fraction (Fig. 1D). This aligns with previous reports by Yancey et al., 2023, who also documented similar patterns of *Microcystis* isolate diversity within the 100 µm fraction during the 2014 bloom in the western basin of Lake Erie.

**Figure 4.**
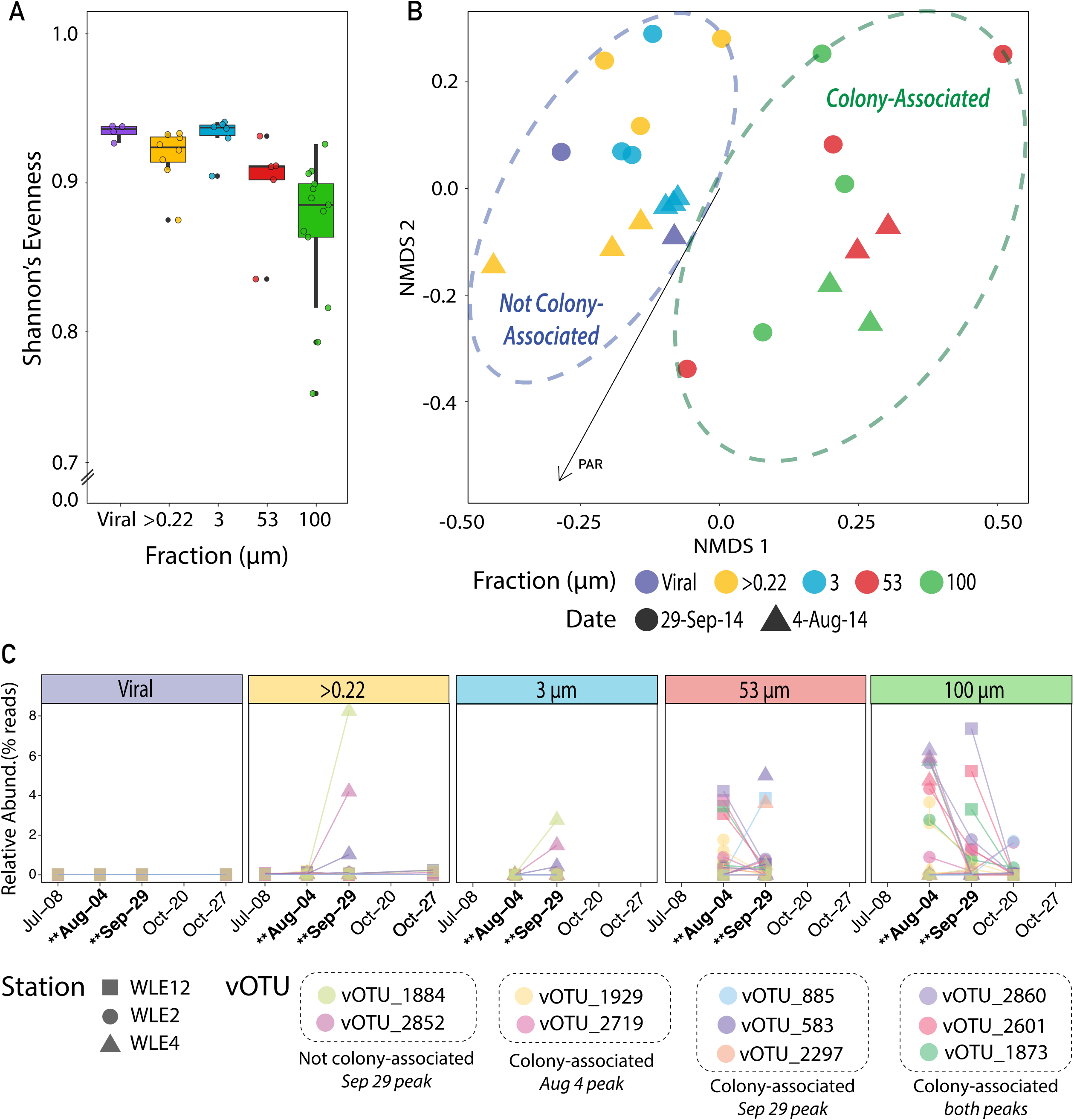
*Microcystis* virus diversity, turnover, and abundant vOTU temporal dynamics through the 2014 Lake Erie bloom season. (A) Diversity of predicted *Microcystis* vOTU assemblage across fractions. (B) NMDS ordination based on Bray-Curtis dissimilarities in distributions of predicted *Microcystis* vOTUs at the cHAB peaks. Point color represents the sampling fraction; shape represents sampling date. Dashed ovals indicate whether a point belongs to the free-living or colony-associated community. The ordinations are overlaid with the gradient of fit between measured environmental parameters and Bray-Curtis dissimilarities. Vector length of the environmental parameter represents the strength of the correlation with the data variation. (C) Temporal relative abundance dynamics of the ten most abundant predicted *Microcystis* vOTUs in terms of fraction of total metagenomic reads mapped to each vOTU per sample. Point shape represents the sampling station and point color represents a specific *Microcystis* virus OTU. Dates of sampled bloom peaks are highlighted in bold with double asterisk.

The observed fraction-specific genotypic variation in *Microcystis* combined with patchy representation of the *Microcystis* MAGs in the colony-associated fractions can be explained by the proliferation of single phylotypes leading to colony formation during bloom development. Such episodes may occur frequently through the Lake Erie bloom, with distinctive phylotypes emerging at each sampling date among 100 µm fraction samples (Fig. 1C). In line with a ‘Kill the Winner’ dynamic,^89^ each such episode may select for a subset of viruses able to infect the dominant phylotype, causing a bottleneck in viral diversity in the colony-associated fractions, which is supported by a corresponding drop in vOTU diversity in the 100 µm fraction (Fig. 4A). In contrast, non-colony associated hosts may sustain infection relationships with more members of the viral “seed bank”,^8,14,23^ thus maintaining a higher overall *Microcystis* vOTU assemblage diversity.

#### Sampling fraction correlates with turnover of vOTUs predicted to infect *Microcystis*

Between-sample variation in the *Microcystis* vOTU assemblage structure significantly correlated with filter fraction (Fig. 4B; PERMANOVA R^2^=0.21, p-value = 0.0001; SI Table 5) and to a lesser extent sampling date (R^2^=0.06, p-value = 0.001). *Microcystis* vOTU assemblages partitioned based on colony-associated versus free-living size fractions (Fig. 4B). This pattern may be explained by the discussed fraction-specific differences in relative abundances of *Microcystis* genotypes (Fig. 1E). Even though free-living and colony-associated communities exist within the same aquatic matrix, they represent distinct ecological contexts allowing for distinct dynamics in host populations,^69^ which we posit also extends to distinct virus-host interactions. These fraction-associated *Microcystis* genotypic differences may have implications for viral host range; viral mediated top down control on *Microcystis* may depend on whether the *Microcystis* host is colonial or single cellular.

Sample location was not significantly correlated with turnover in the *Microcystis* vOTU assemblage, (R^2^=0.08, p-value = 0.30). In understanding top down controls on *Microcystis*, factors that underlie the dynamics of a single *Microcystis* virus population (let alone a single gene) will not reflect the dynamics of the total assemblage of *Microcystis* viruses. Of the environmental variables tested, only photoactive radiation (PAR) was significantly correlated with variation in *Microcystis* vOTU turnover (Fig. 4B; R^2^= 0.06, p-value = 0.0014; SI Table 6).

Given that *Microcystis* growth and physiology are influenced by variations in light intensity and duration,^88^ light may indirectly affect the viruses via effects of PAR on the host population. This trend may also suggest light availability as a driver of *Microcystis* infection during high bloom densities.

#### High turnover of abundant *Microcystis* virus populations

We next sought to better understand the seasonal fluctuation and host ranges of the numerically important *Microcystis* vOTUs in the 2014 Lake Erie cHAB. We identified the ten *Microcystis* vOTUs most abundant across the entire sample set (SI Table 2). Notably, *Microcystis* phage Ma-LMM01, whose gp91 gene has been used as the sole proxy for quantifying *Microcystis* viruses in some studies,^39,51,83,94^ was not among these abundant vOTUs. When relatively assessed per fraction, the dominant *Microcystis* vOTUs in the cellular fractions were never abundant in the viral fraction (Fig. 1C). When they peaked, they were most abundant at the bloom peaks (Fig. 4C). They peaked in abundance in either the colony-associated or unassociated fractions during bloom peaks on either Aug 4 or Sept 29, with only three of the 10 vOTUs peaking at both dates (Fig. 4C).

The sporadic peaks of these abundant *Microcystis* vOTUs are consistent with previous descriptions of viral seed bank, i.e., Bank dynamics,^8,23^ whereby only a few *Microcystis* vOTUs are abundant at any given moment throughout the seasons—and those that do rise to high abundance do so in the cellular fractions where active infections presumably dominate. These changes in viral community composition over time are likely influenced by factors such as shifts in host availability, environmental conditions, and the viral populations’ specific interactions with *Microcystis* and other host populations. Indeed, even at the coarse level of oligotypes, different *Microcystis* genotypes were present at the two bloom peaks.^5^ Of the 10 most abundant *Microcystis* viral populations identified, 80% were primarily found in either the 53 µm or 100 µm fraction, which represents the *Microcystis* colony-associated viral community. As host cells within colonies are tightly-packed, the opportunity to infect and increase in abundance is more likely to occur here than in those viral populations that persist in the free-living microbial community, where host interactions may be scarce. Alternatively, colonies may also provide defense against infection.^90^ Understanding the turnover and dynamics of these viral populations is essential for comprehending their roles in regulating harmful algal blooms and the broader dynamics of aquatic ecosystems in Lake Erie.

## Conclusions and Outlook

Previous research has established that the two 2014 bloom peaks represented different *Microcystis* genotypes,^92^ including the transition from toxin-producing to non-toxin producing genotypes. This was hypothesized to be in part due to the low ammonium and nitrate availability in September relative to August coupled with the nitrogen-rich nature of microcystin metabolites.^13,92^ Our work suggests viral activity as an additional control on *Microcystis* strain succession in the 2014 bloom. Distinct assemblages of *Microcystis* vOTUs were predicted to infect the *Microcystis* genotypes at the two bloom peaks, suggesting strain specificity among predicted *Microcystis* viruses. Strain-specific viral predation and the diversity and ecological dynamics of *Microcystis* populations are inextricably connected, likely influencing one another over the course of the bloom.

The patterns uncovered in this work describing the spatial and temporal dynamics of *Microcystis* viruses in the 2014 Lake Erie cHAB help to reveal the intricate dynamics of cyanobacterial blooms and their ecological implications. Additional molecular techniques, such as metatranscriptomics, metaproteomics, and metabolomics, will help to identify viral gene and protein expression patterns and to identify relationships between viral activity and community-level metabolite (including toxin) production through the blooms. These omics approaches can also shed light on viral strategies for manipulating host metabolism, toxin regulation, and avoiding host antiviral strategies within cHABs. Moreover, longitudinal studies encompassing multiple bloom seasons and locations will contribute valuable insights into the temporal and spatial dynamics of *Microcystis* vOTUs. Understanding connections between the *Microcystis* viruses, the total virus community, environmental parameters, and overall bloom progression is essential for unraveling the complex dynamics of these cHABs.

## Materials and Methods

### Field sampling and collection

Field sites were sampled with the joint NOAA Great Lakes Environmental Research Laboratory / University of Michigan Cooperative Institute for Great Lakes Research weekly sampling program for Lake Erie. In 2014, three sites were sampled bi-monthly in June then weekly from July through October. Bloom stages were determined by phycocyanin fluorescence and relative abundance. Water chemistry measurements are detailed in Berry et al., 2017. Metagenomic data were generated from samples collected from three regularly sampled stations (WE2, near the mouth of Maumee River, 41° 45.743’ N, 83°19.874’ W; WE4, offshore towards the center of the western basin, 41°49.595’ N, 83°11.698’ W; and WE12, adjacent to the water intake crib for the city of Toledo, 41°42.535’ N, 83°14.989’ W). All samples were collected upon arriving on-station using a peristaltic pump to obtain 20 L of water from 0.1 m below the surface. Water was filtered onto 100 μm polycarbonate filters. This size was used to concentrate *Microcystis* colonies retained on the filter while excluding smaller particles. Previous work has shown that in Lake Erie the >100 μm assemblage comprised over 90% of all *Microcystis* cells in the water column.^11^ The filtered water was subsequently passed through a 53 µm and 3 µm polycarbonate filter to collect smaller colonies and large single-celled microbes, including *Microcystis*, whose cell sizes range from 1.7 to 7 µm in diameter.^91^ Whole water was passed through a 0.22 µm filter to collect the total cellular microbial community; community structure of whole community fractions have been shown to be dominated by single cells.^75^ To enrich for viruses 10 g/L iron chloride stock solution (0.966 g FeCl_3_-6H_2_O in 20 mL 0.02 µm-filtered autoclaved MilliQ-H_2_O) was added to the <0.22 µm (“viral”) fraction.^65^ The flocculant incubated overnight to maximize virus recovery before being passed through a 0.45 µm 142 mm Millipore Express Plus filter and stored at 4°C.

### DNA extraction and sequencing of hosts and viruses

DNA was extracted from samples using the DNeasy Mini Kit (QIAGEN) according to the manufacturer’s instructions. Shotgun sequencing of DNA was performed on the Illumina® HiSeq™ platform (2000 PE 100, Illumina, Inc., San Diego, CA, USA) at the University of Michigan DNA Sequencing Core.

### Host assembly and binning

BBDuk (https://sourceforge.net/projects/bbduk/) was used for read quality trimming, length trimming (anything less than 100bp), identifying and removing contaminated sequences against univec, and evaluating denoised reads using FastQC (http://www.bioinformatics.bbsrc.ac.uk/projects/fastqc/). The reads were dereplicated using BBnorm (https://sourceforge.net/projects/bbnorm/). Reads of all 36 samples were assembled on a per-sample basis into contigs using Megahit with metasensitive parameters.^44^ Following contig assembly, Centrifuge was used to taxonomically classify reads to the species level, or lowest resolution available.^38^ Quality trimmed raw reads were mapped to each individual assembly using bwa (https://sourceforge.net/projects/bio-bwa/) with bwa-mem on default settings.

SAMtools was used to convert, sort, and index compressed BAM files.^45^ Quality trimmed reads were competitively mapped to MAGs using a pileup shell script provided by BBtools (https://sourceforge.net/projects/bbtools/). Automated binning was performed on the contigs using Concoct on default settings to generate metagenome assembled genomes (MAGs).^1^ Bins with >50% completeness (completeness statistics inferred from a CheckM),^63^ >10% contamination, and <75% strain heterogeneity were manually refined in Anvi’o based on differential coverage and contamination. The Anvi’o platform v2.3.0 was used to manually refine the unique MAGs identified through Concoct and Vizbin by evaluating differential coverage patterns across the samples.^16,42^ Multiple rounds of Anvi’o refinement were performed to curate bins until they passed the aforementioned thresholds (SI Table 7).

### Viral population identification and taxonomic assignment

CheckV v0.7.0,^58^ VIBRANT v1.2.1,^36^ VirFinder v1.1,^68^ VirSorter v1.0.6,^71^ and VirSorter2 v2.1 were used to predict presumed viral contigs.^19^ Contigs from each tool were preserved according to recently established rules for viral contig identification.^21^ Additionally, only contigs >3kb were kept from the viral prediction tools and used to identify viral populations. Viruses were then binned using vRhyme default settings to create a collection of viral bins and high-quality unbinned contigs for population clustering.^35^ Viral bins and unbinned contigs were clustered (stampede-clustergenomes)^71^ according to previously established standards defining viral populations.^72^ Contigs sharing an average nucleotide identity (ANI) of 95% across 85% of the contig length were clustered and the longest sequence of each cluster was considered the representative for a cluster, referred to as a viral operational taxonomic unit (vOTU). Taxonomy of viral populations from the two Lake Erie bloom peaks was estimated using the Phage Taxonomy Tool approach (PTT).^37^

### Viral community read mapping, quantification and alpha/beta diversity

Filtered and trimmed reads were assembled from the same sample and quantified using Samtools v1.11.^45^ These reads were then competitively mapped to all contigs within a given sample using Bowtie2.^43^ Reads mapped to trimmed viral sequences were quantified using BLAST v2.9.0 to align trimmed viral sequences to the contigs and then using FeatureCounts from the Subread package to quantify reads overlapping this region.^46,47,59^ The viral reads for each sample were downsampled by 1,000,000 reads for alpha diversity analyses using seqtk v1-3 (http://github.com/Ih3/seqtk). Only viral contigs with reads covering at least 3 kb of their length were extracted and included in diversity analyses. Alpha diversity measures were calculated using the vegan v2.5-2 package in R v4.0.2 based on downsampled reads.

Transcripts per million (TPM) was determined for each viral population and used to calculate the Bray-Curtis distance between samples in R using the vegan package and then NMDS ordination was performed. PERMANOVA using the *adonis* function in vegan was used to test the effects of sampling location, sampling date, sampling fraction as well as effects of environmental parameters on the full viral community structure and metabolic potential. Viral contig information and NCBI accession numbers can be found in SI Table 8.

### Viral metabolic potential analyses

KEGG and Pfam databases were accessed to assign viral contig gene metabolic annotations using Distilled and Refined Annotation of Metabolism (DRAM)^76^ following the generation of open reading frames (ORFs) using Prodigal.^29^ FeatureCounts from the Subread package was then used to calculate read coverages of the ORFs generated by DRAM.^46,47^ Bray Curtis distances between samples were calculated using the vegdist function followed by an NMDS ordination with the vegan package in R. Water quality parameters were then applied to a PERMANOVA model to evaluate their effects on the abundance of protein families (Pfams). Parameters were considered significantly correlated with a p-value ≤ 0.05.

### Virus-host infection prediction network

To better understand potential phage infections associated with bacterial hosts in the bloom, we applied VHIP (v.1.0), a gradient-boosted machine learning model informed by signals of coevolution embedded in sequences between viruses and hosts.^3^ All possible virus-host combinations were considered initially. For network visualization, only viral sequences larger than 10kb and prediction scores higher than 0.93 were considered. (viral sequences >10kb and a >93% probability of infection) were plotted using Gephi 0.9.0 software.

## Availability of data and materials

Read data sets are publicly available in NCBI under SRA numbers: SRX4099271 to SRX4099280, SRX4099286 to SRX4099288, and SRX4099293 to SRX4099300 in BioProject no. PRJNA464361 and SRX24032252 to SRX24032266 in BioProject no. PRJN988094. All metagenomic assemblies used to generate host MAGs and identify viral contigs are available under BioSample numbers: SAMN36000421 to SAMN36000456 in BioProject no. PRJN988094. The scripts used here, including viral identification tools, taxonomy assignment, relative abundance calculations, are freely available at https://github.com/DuhaimeLab/Tracking_Ma_viruses_and_infection_dynamics_across_distinct_Ma_bloom_phases.

## Acknowledgements

We would like to thank Michelle Berry for early assistance with viral sample collection and processing and Lizy Michaelson for assistance verifying and publishing protocols in protocols.io. We would also like to thank Derek Smith for his efforts and manual curation of metagenomic host MAGs used for this study. We acknowledge the members of the Duhaime Lab for years of constructive feedback on the research and writing that went into this work. We are grateful to the NOAA Great Lakes Environmental Research Lab and the Cooperative Institute for Great Lakes Research logistical team and field crew for allowing us to sample with their HAB monitoring efforts and for assisting in field measurements.

## References

1) Alneberg, J., Bjarnason, B. S., de Bruijn, I., Schirmer, M., Quick, J., Ijaz, U. Z., Loman, N. J., Andersson, A. F., & Quince, C. (2013). *CONCOCT: Clustering cONtigs on COverage and ComposiTion* (arXiv:1312.4038). arXiv. 10.48550/arXiv.1312.4038

2 Anantharaman, K., Duhaime, M. B., Breier, J. A., Wendt, K. A., Toner, B. M., & Dick, G. J. (2014). Sulfur Oxidation Genes in Diverse Deep-Sea Viruses. Science, 344(6185), 757–760. 10.1126/science.1252229

3 Bastien, E. G., Cable, R. N., Zaman, L., Batterbee, C., Wing, A. J., & Duhaime, M. B. (2023). Virus-Host Interactions Predictor (VHIP): Machine learning approach to resolve microbial virus-host interaction networks (p. 2023.11.03.565433). bioRxiv. 10.1101/2023.11.03.565433

4) Berry, M. A., Davis, T. W., Cory, R. M., Duhaime, M. B., Johengen, T. H., Kling, G. W., Marino, J. A., Den Uyl, P. A., Gossiaux, D., Dick, G. J., & Denef, V. J. (2017). Cyanobacterial harmful algal blooms are a biological disturbance to Western Lake Erie bacterial communities. 10.1111/1462-2920.13640

5 Berry, M. A., White, J. D., Davis, T. W., Jain, S., Johengen, T. H., Dick, G. J., Sarnelle, O., & Denef, V. J. (2017b). Are Oligotypes Meaningful Ecological and Phylogenetic Units? A Case Study of *Microcystis* in Freshwater Lakes. Frontiers in Microbiology, 08, 365–365. 10.3389/fmicb.2017.00365

6 Bolay, P., Muro-Pastor, M. I., Florencio, F. J., & Klähn, S. (2018). The Distinctive Regulation of Cyanobacterial Glutamine Synthetase. *Life (Basel*, Switzerland), 8(4), 52. 10.3390/life8040052

7 Breitbart, M. (2012). Marine viruses: Truth or dare. Annual Review of Marine Science, 4, 425–448. 10.1146/annurev-marine-120709-142805

8 Breitbart, M., & Rohwer, F. (2005). Here a virus, there a virus, everywhere the same virus? Trends in Microbiology, 13(6), 278–284. 10.1016/j.tim.2005.04.003

9 Burstein, D., Sun, C. L., Brown, C. T., Sharon, I., Anantharaman, K., Probst, A. J., Thomas, B. C., & Banfield, J. F. (2016). Major bacterial lineages are essentially devoid of CRISPR-Cas viral defence systems. Nature Communications, 7(1), Article 1. 10.1038/ncomms10613

10 Cai, R., Li, D., Lin, W., Qin, W., Pan, L., Wang, F., Qian, M., Liu, W., Zhou, Q., Zhou, C., & Tong, Y. (2022). Genome sequence of the novel freshwater *Microcystis* cyanophage Mwe-Yong1112-1. Archives of Virology, 167, 1–6. 10.1007/s00705-022-05542-3

11 Chaffin, J. D., Bridgeman, T. B., Heckathorn, S. A., & Mishra, S. (2011). Assessment of *Microcystis* growth rate potential and nutrient status across a trophic gradient in western Lake Erie. Journal of Great Lakes Research, 37(1), 92–100. 10.1016/j.jglr.2010.11.016

12 Correa, A. M. S., Howard-Varona, C., Coy, S. R., Buchan, A., Sullivan, M. B., & Weitz, J. S. (2021). Revisiting the rules of life for viruses of microorganisms. Nature Reviews Microbiology, 19(8), Article 8. 10.1038/s41579-021-00530-x

13 Cory, R. M., Davis, T. W., Dick, G. J., Johengen, T., Denef, V. J., Berry, M. A., Page, S. E., Watson, S. B., Yuhas, K., & Kling, G. W. (2016). Seasonal Dynamics in Dissolved Organic Matter, Hydrogen Peroxide, and Cyanobacterial Blooms in Lake Erie. Frontiers in Marine Science, 3. https://www.frontiersin.org/articles/10.3389/fmars.2016.00054

14 Dart, E., Fuhrman, J. A., & Ahlgren, N. A. (2023). Diverse Marine T4-like Cyanophage Communities Are Primarily Comprised of Low-Abundance Species Including Species with Distinct Seasonal, Persistent, Occasional, or Sporadic Dynamics. Viruses, 15(2), Article 2. 10.3390/v15020581

15 Enav, H., Kirzner, S., Lindell, D., Mandel-Gutfreund, Y., & Béjà, O. (2018). Adaptation to sub-optimal hosts is a driver of viral diversification in the ocean. Nature Communications, 9, 4698. 10.1038/s41467-018-07164-3

16 Eren, A. M., Kiefl, E., Shaiber, A., Veseli, I., Miller, S. E., Schechter, M. S., Fink, I., Pan, J. N., Yousef, M., Fogarty, E. C., Trigodet, F., Watson, A. R., Esen, Ö. C., Moore, R. M., Clayssen, Q., Lee, M. D., Kivenson, V., Graham, E. D., Merrill, B. D., … Willis, A. D. (2021). Community-led, integrated, reproducible multi-omics with anvi’o. Nature Microbiology, 6(1), Article 1. 10.1038/s41564-020-00834-3

17) Fuhrman. (1999). *Marine viruses and their biogeochemical and ecological effects | Natur*e. https://www.nature.com/articles/21119

18) Garcia-Villegas, M., De La Vega, M., Galindo, J., Segura, M., Buckingham, R., Guarneros, G. (1991). Peptidyl-tRNA hydrolase is involved in lambda inhibition of host protein synthesis. Retrieved January 10, 2024, from https://www.embopress.org/doi/epdf/10.1002/j.1460-2075.1991.tb04919.x

19 Guo, J., Bolduc, B., Zayed, A. A., Varsani, A., Dominguez-Huerta, G., Delmont, T. O., Pratama, A. A., Gazitúa, M. C., Vik, D., Sullivan, M. B., & Roux, S. (2021). VirSorter2: A multi-classifier, expert-guided approach to detect diverse DNA and RNA viruses. Microbiome, 9(1), 37. 10.1186/s40168-020-00990-y

20 Harke, M. J., Steffen, M. M., Gobler, C. J., Otten, T. G., Wilhelm, S. W., Wood, S. A., & Paerl, H. W. (2016). A review of the global ecology, genomics, and biogeography of the toxic cyanobacterium, Microcystis spp. Harmful Algae, 54, 4–20. 10.1016/j.hal.2015.12.007

21 Hegarty, B., Dai, Z., Raskin, L., Pinto, A., Wigginton, K., & Duhaime, M. (2022). A snapshot of the global drinking water virome: Diversity and metabolic potential vary with residual disinfectant use. Water Research, 218, 118484. 10.1016/j.watres.2022.118484

22 Herr, C. Q., & Hausinger, R. P. (2018). Amazing Diversity in Biochemical Roles of Fe(II)/2-Oxoglutarate Oxygenases. Trends in Biochemical Sciences, 43(7), 517–532. 10.1016/j.tibs.2018.04.002

23 Hevroni, G., Flores-Uribe, J., Béjà, O., & Philosof, A. (2020). Rising through the Ranks: Seasonal and Diel Patterns of Marine Viruses. bioRxiv, 2020.05.07.082883. 10.1101/2020.05.07.082883

24 Howard-Varona, C., Lindback, M. M., Bastien, G. E., Solonenko, N., Zayed, A. A., Jang, H., Andreopoulos, B., Brewer, H. M., Glavina del Rio, T., Adkins, J. N., Paul, S., Sullivan, M. B., & Duhaime, M. B. (2020). Phage-specific metabolic reprogramming of virocells. The ISME Journal, 14(4), Article 4. 10.1038/s41396-019-0580-z

25 Huisman, J., Codd, G. A., Paerl, H. W., Ibelings, B. W., Verspagen, J. M. H., & Visser, P. M. (2018). Cyanobacterial blooms. Nature Reviews Microbiology, 16(8), 471–483. 10.1038/s41579-018-0040-1

26 Hunt, D. E., David, L. A., Gevers, D., Preheim, S. P., Alm, E. J., & Polz, M. F. (2008). Resource Partitioning and Sympatric Differentiation Among Closely Related Bacterioplankton. Science, 320(5879), 1081–1085. 10.1126/science.1157890

27 Hurwitz, B. L., Hallam, S. J., & Sullivan, M. B. (2013). Metabolic reprogramming by viruses in the sunlit and dark ocean. Genome Biology, 14(11), R123. 10.1186/gb-2013-14-11-r123

28 Hwang, Y., Roux, S., Coclet, C., Krause, S. J. E., & Girguis, P. R. (2023). Viruses interact with hosts that span distantly related microbial domains in dense hydrothermal mats. Nature Microbiology, 8(5), Article 5. 10.1038/s41564-023-01347-5

29 Hyatt, D., Chen, G.-L., LoCascio, P. F., Land, M. L., Larimer, F. W., & Hauser, L. J. (2010). Prodigal: Prokaryotic gene recognition and translation initiation site identification. BMC Bioinformatics, 11(1), 119. 10.1186/1471-2105-11-119

30) Jain, et al. (2018). High throughput ANI analysis of 90K prokaryotic genomes reveals clear species boundaries | Nature Communications. https://www.nature.com/articles/s41467-018-07641-9

31 Jantaro, S., Mäenpää, P., Mulo, P., & Incharoensakdi, A. (2003). Content and biosynthesis of polyamines in salt and osmotically stressed cells of Synechocystis sp. PCC 6803. FEMS Microbiology Letters, 228(1), 129–135. 10.1016/S0378-1097(03)00747-X

32 Jia, B., Jia, X., Kim, K. H., & Jeon, C. O. (2017). Integrative view of 2-oxoglutarate/Fe(II)-dependent oxygenase diversity and functions in bacteria. Biochimica et Biophysica Acta (BBA) - General Subjects, 1861(2), 323–334. 10.1016/j.bbagen.2016.12.001

33 Jiang, X., Ha, C., Lee, S., Kwon, J., Cho, H., Gorham, T., & Lee, J. (2019). Characterization of Cyanophages in Lake Erie: Interaction Mechanisms and Structural Damage of Toxic Cyanobacteria. Toxins, 11(8), 444. 10.3390/toxins11080444

34 Kehr, J.C., & Dittmann, E. (2015). Biosynthesis and function of extracellular glycans in cyanobacteria. *Life (Basel*, Switzerland), 5(1), 164–180. 10.3390/life5010164

35) Kieft, K., Adams, A., Salamzade, R., Kalan, L., Anantharaman, K. (2022). vRhyme enables binning of viral genomes from metagenomes | Nucleic Acids Research | Oxford Academic. https://academic.oup.com/nar/article/50/14/e83/6584432

36 Kieft, K., Zhou, Z., & Anantharaman, K. (2019). VIBRANT: Automated recovery, annotation and curation of microbial viruses, and evaluation of virome function from genomic sequences. bioRxiv, 855387. 10.1101/855387

37 Kieft, K., Zhou, Z., Anderson, R. E., Buchan, A., Campbell, B. J., Hallam, S. J., Hess, M., Sullivan, M. B., Walsh, D. A., Roux, S., & Anantharaman, K. (2021). Ecology of inorganic sulfur auxiliary metabolism in widespread bacteriophages. Nature Communications, 12(1), Article 1. 10.1038/s41467-021-23698-5

38 Kim, J.-G., Park, S.-J., Sinninghe Damsté, J. S., Schouten, S., Rijpstra, W. I. C., Jung, M.-Y., Kim, S.-J., Gwak, J.-H., Hong, H., Si, O.-J., Lee, S., Madsen, E. L., & Rhee, S.-K. (2016). Hydrogen peroxide detoxification is a key mechanism for growth of ammonia-oxidizing archaea. Proceedings of the National Academy of Sciences of the United States of America, 113(28), 7888–7893. 10.1073/pnas.1605501113

39 Kimura, S., Sako, Y., & Yoshida, T. (2012). Rapid *Microcystis* cyanophage gene diversification revealed by long- and short-term genetic analyses of the tail sheath gene in a natural pond. Applied and Environmental Microbiology, 79(8), 2789–2795. 10.1128/AEM.03751-12

40 Koskella, B., & Brockhurst, M. A. (2014). Bacteria-phage coevolution as a driver of ecological and evolutionary processes in microbial communities. FEMS Microbiology Reviews, 38(5), 916–931. 10.1111/1574-6976.12072

41 Krzywinski, M., Schein, J., Birol, I., Connors, J., Gascoyne, R., Horsman, D., Jones, S. J., & Marra, M. A. (2009). Circos: An information aesthetic for comparative genomics. Genome Research, 19(9), 1639–1645. 10.1101/gr.092759.109

42) Laczny, C. C., Sternal, T., Plugaru, V., Gawron, P., Atashpendar, A., Margossian, H. H., Coronado, S., der Maaten, L. van, Vlassis, N., & Wilmes, P. (2015). VizBin—An application for reference-independent visualization and human-augmented binning of metagenomic data. Microbiome, 3(1), 1. 10.1186/s40168-014-0066-1

43 Langmead, B., & Salzberg, S. L. (2012). Fast gapped-read alignment with Bowtie 2. Nature Methods, 9(4), Article 4. 10.1038/nmeth.1923

44 Li, D., Liu, C.-M., Luo, R., Sadakane, K., & Lam, T.-W. (2015). MEGAHIT: An ultra-fast single-node solution for large and complex metagenomics assembly via succinct de Bruijn graph. Bioinformatics, 31(10), 1674–1676. 10.1093/bioinformatics/btv033

45 Li, H., Handsaker, B., Wysoker, A., Fennell, T., Ruan, J., Homer, N., Marth, G., Abecasis, G., Durbin, R., & 1000 Genome Project Data Processing Subgroup. (2009). The Sequence Alignment/Map format and SAMtools. Bioinformatics (Oxford, England), 25(16), 2078–2079. 10.1093/bioinformatics/btp352

46 Liao, Y., Smyth, G. K., & Shi, W. (2013). The Subread aligner: Fast, accurate and scalable read mapping by seed-and-vote. Nucleic Acids Research, 41(10), e108. 10.1093/nar/gkt214

47 Liao, Y., Smyth, G. K., & Shi, W. (2014). featureCounts: An efficient general purpose program for assigning sequence reads to genomic features. *Bioinformatics (Oxford*, England), 30(7), 923–930. 10.1093/bioinformatics/btt656

48 Lindell, D., Sullivan, M. B., Johnson, Z. I., Tolonen, A. C., Rohwer, F., & Chisholm, S. W. (2004). Transfer of photosynthesis genes to and from Prochlorococcus viruses. Proceedings of the National Academy of Sciences of the United States of America, 101(30), 11013–11018. 10.1073/pnas.0401526101

49 Luque, A., & Silveira, C. B. (2020). Quantification of Lysogeny Caused by Phage Coinfections in Microbial Communities from Biophysical Principles. mSystems, 5(5), 10.1128/msystems.00353-20.

50 Malki, K., Kula, A., Bruder, K., Sible, E., Hatzopoulos, T., Steidel, S., Watkins, S. C., & Putonti, C. (2015). Bacteriophages isolated from Lake Michigan demonstrate broad host-range across several bacterial phyla. Virology Journal, 12(1), 164. 10.1186/s12985-015-0395-0

51 Mankiewicz-Boczek, J., Jaskulska, A., Pawełczyk, J., Gągała, I., Serwecińska, L., & Dziadek, J. (2016). Cyanophages Infection of *Microcystis* Bloom in Lowland Dam Reservoir of Sulejów, Poland. Microbial Ecology, 71(2), 315–325. 10.1007/s00248-015-0677-5

52 McDaniel, L. D., Young, E., Delaney, J., Ruhnau, F., Ritchie, K. B., & Paul, J. H. (2010). High frequency of horizontal gene transfer in the oceans. Science (New York, N.Y.), 330(6000), 50. 10.1126/science.1192243

53 McKindles, K. M., Manes, M. A., DeMarco, J. R., McClure, A., McKay, R. M., Davis, T. W., & Bullerjahn, G. S. (2020). Dissolved Microcystin Release Coincident with Lysis of a Bloom Dominated by *Microcystis* spp. In Western Lake Erie Attributed to a Novel Cyanophage. Applied and Environmental Microbiology. 10.1128/AEM.01397-20

54) Michalek, et al. (2013). Record-setting algal bloom in Lake Erie caused by agricultural and meteorological trends consistent with expected future conditions | PNAS. https://www-pnas-org.proxy.lib.umich.edu/doi/10.1073/pnas.1216006110

55 Morimoto, D., Tominaga, K., Nishimura, Y., Yoshida, N., Kimura, S., Sako, Y., & Yoshida, T. (2019). Co-occurrence of broad and narrow host-range viruses infecting the toxic bloom-forming cyanobacterium *Microcystis aeruginosa*. *Applied and Environmental Microbiology*, AEM.01170–19. 10.1128/AEM.01170-19

56 Morimoto, D., Yoshida, N., Sasaki, A., Nakagawa, S., Sako, Y., & Yoshida, T. (2023). Ecological Dynamics of Broad- and Narrow-Host-Range Viruses Infecting the Bloom-Forming Toxic Cyanobacterium *Microcystis aeruginosa*. Applied and Environmental Microbiology, 89(2), e02111–22. 10.1128/aem.02111-22

57 Morohoshi, T., Maruo, T., Shirai, Y., Kato, J., Ikeda, T., Takiguchi, N., Ohtake, H., & Kuroda, A. (2002). Accumulation of inorganic polyphosphate in phoU mutants of Escherichia coli and Synechocystis sp. Strain PCC6803. Applied and Environmental Microbiology, 68(8), 4107–4110. 10.1128/AEM.68.8.4107-4110.2002

58 Nayfach, S., Camargo, A. P., Schulz, F., Eloe-Fadrosh, E., Roux, S., & Kyrpides, N. C. (2021). CheckV assesses the quality and completeness of metagenome-assembled viral genomes. Nature Biotechnology, 39(5), 578–585. 10.1038/s41587-020-00774-7

59 NCBI. (2019, April 2). *BLAST+ v2.9.0*. NCBI Insights. https://ncbiinsights.ncbi.nlm.nih.gov/2019/04/02/blast-2-9-0-now-available-with-enhanced-support-for-new-database-format-and-improved-performance/

60) O’neil, J.M., Davis, T.W., Buford, M.A. & Gobler, C.J. (2012). The rise of harmful cyanobacteria blooms: The potential roles of eutrophication and climate change— ScienceDirect. https://www-sciencedirect-com.proxy.lib.umich.edu/science/article/pii/S1568988311001557

61) Paerl, H. W. and Huisman, J. (2008). Blooms Like It Hot | Science. https://www.science.org/doi/10.1126/science.1155398

62 Paerl, H. W. and Huisman, J. (2009). Climate change: A catalyst for global expansion of harmful cyanobacterial blooms. Environmental Microbiology Reports, 1(1), 27–37. 10.1111/j.1758-2229.2008.00004.x

63 Parks, D. H., Imelfort, M., Skennerton, C. T., Hugenholtz, P., & Tyson, G. W. (2015). CheckM: Assessing the quality of microbial genomes recovered from isolates, single cells, and metagenomes. Genome Research, 25(7), 1043–1055. 10.1101/gr.186072.114

64 Pérez-Carrascal, O. M., Terrat, Y., Giani, A., Fortin, N., Greer, C. W., Tromas, N., & Shapiro, B. J. (2019). Coherence of *Microcystis* species revealed through population genomics. The ISME Journal, 13(12), 2887–2900. 10.1038/s41396-019-0481-1

65 Poulos, B., John, S., Sullivan, M.B. (2017). Iron Chloride Flocculation of Bacteriophages from Seawater. Bacteriophages, 1681. https://link-springer-com.proxy.lib.umich.edu/protocol/10.1007/978-1-4939-7343-9_4

66 Pound, H. L., & Wilhelm, S. W. (2020). Tracing the active genetic diversity of *Microcystis* and *Microcystis* phage through a temporal survey of Taihu. PLoS ONE, 15(12), e0244482. 10.1371/journal.pone.0244482

67 Preece, E. P., Hardy, F. J., Moore, B. C., & Bryan, M. (2017). A review of microcystin detections in Estuarine and Marine waters: Environmental implications and human health risk. Harmful Algae, 61, 31–45. 10.1016/j.hal.2016.11.006

68 Ren, J., Ahlgren, N. A., Lu, Y. Y., Fuhrman, J. A., & Sun, F. (2017). VirFinder: A novel k-mer based tool for identifying viral sequences from assembled metagenomic data. Microbiome, 5(1), 69. 10.1186/s40168-017-0283-5

69 Rosenberg, D., Haber, M., Goldford, J., Lalzar, M., Aharonovich, D., Al-Ashhab, A., Lehahn, Y., Segrè, D., Steindler, L., & Sher, D. (2021). Particle-associated and free-living bacterial communities in an oligotrophic sea are affected by different environmental factors. Environmental Microbiology, 23(8), 4295–4308. 10.1111/1462-2920.15611

70 Rosenwasser, S., Ziv, C., Creveld, S. G. van, & Vardi, A. (2016). Virocell Metabolism: Metabolic Innovations During Host-Virus Interactions in the Ocean. Trends in Microbiology, 24(10), 821–832. 10.1016/j.tim.2016.06.006

71) Roux and Bolduc. (2015). MAVERICLab / stampede-clustergenomes—Bitbucket. https://bitbucket.org/MAVERICLab/stampede-clustergenomes/src/master/

72 Roux, S., Adriaenssens, E. M., Dutilh, B. E., Koonin, E. V., Kropinski, A. M., Krupovic, M., Kuhn, J. H., Lavigne, R., Brister, J. R., Varsani, A., Amid, C., Aziz, R. K., Bordenstein, S. R., Bork, P., Breitbart, M., Cochrane, G. R., Daly, R. A., Desnues, C., Duhaime, M. B., … Eloe-Fadrosh, E. A. (2019). Minimum Information about an Uncultivated Virus Genome (MIUViG). Nature Biotechnology, 37(1), Article 1. 10.1038/nbt.4306

73 Roux, S., Enault, F., Hurwitz, B. L., & Sullivan, M. B. (2016). VirSorter: Mining viral signal from microbial genomic data. PeerJ, 3, e985. 10.7717/peerj.985

74) Rozon and Short. (2013). Complex seasonality observed amongst diverse phytoplankton viruses in the Bay of Quinte, an embayment of Lake Ontario—Rozon—2013— Freshwater Biology—Wiley Online Library. https://onlinelibrary-wiley-com.proxy.lib.umich.edu/doi/full/10.1111/fwb.12241

75 Schmidt, K. C., Jackrel, S. L., Smith, D. J., Dick, G. J., & Denef, V. J. (2020). Genotype and host microbiome alter competitive interactions between *Microcystis aeruginosa* and Chlorella sorokiniana. Harmful Algae, 99, 101939. 10.1016/j.hal.2020.101939

76) Shaffer, et al. (2020). DRAM for distilling microbial metabolism to automate the curation of microbiome function | Nucleic Acids Research | Oxford Academic. https://academic.oup.com/nar/article/48/16/8883/5884738

77 Smith, D. J., Berry, M. A., Cory, R. M., Johengen, T. H., Kling, G. W., Davis, T. W., & Dick, G. J. (2022). Heterotrophic Bacteria Dominate Catalase Expression during *Microcystis* Blooms. Applied and Environmental Microbiology, 88(14), e0254421. 10.1128/aem.02544-21

78 Smith, D. J., Tan, J. Y., Powers, M. A., Lin, X. N., Davis, T. W., & Dick, G. J. (2021). Individual *Microcystis* colonies harbour distinct bacterial communities that differ by *Microcystis* oligotype and with time. Environmental Microbiology, 23(6), 3020–3036. 10.1111/1462-2920.15514

79 Soucy, S. M., Huang, J., & Gogarten, J. P. (2015). Horizontal gene transfer: Building the web of life. Nature Reviews Genetics, 16(8), Article 8. 10.1038/nrg3962

80 Steffen, M. M., Davis, T. W., McKay, R. M., Bullerjahn, G. S., Krausfeldt, L. E., Stough, J. M. A., Neitzey, M. L., Gilbert, N. E., Boyer, G. L., Johengen, T. H., Gossiaux, D. C., Burtner, A. M., Rowe, M., Dick, G. J., Meyer, K., Levy, S., Boone, B., Wynne, T., Zimba, P. V., … Wilhelm, S. W. (2017). Ecophysiological examination of the Lake Erie *Microcystis* bloom in 2014: Linkages between biology and the water supply shutdown of Toledo, Ohio. Environmental Science.

81 Sullivan, M. J., Petty, N. K., & Beatson, S. A. (2011). Easyfig: A genome comparison visualizer. *Bioinformatics (Oxford*, England), 27(7), 1009–1010. 10.1093/bioinformatics/btr039

82 Suttle, C. A. (2007). Marine viruses—Major players in the global ecosystem. Nature Reviews. Microbiology, 5(10), 801–812. 10.1038/nrmicro1750

83 Takashima, Y., Yoshida, T., Yoshida, M., Shirai, Y., Tomaru, Y., Takao, Y., Hiroishi, S., & Nagasaki, K. (2007). Development and Application of Quantitative Detection of Cyanophages Phylogenetically Related to Cyanophage Ma-LMM01 Infecting *Microcystis aeruginosa* in Fresh Water. Microbes and Environments, 22(3), 207–213. 10.1264/jsme2.22.207

84 Visser, P. M., Verspagen, J. M. H., Sandrini, G., Stal, L. J., Matthijs, H. C. P., Davis, T. W., Paerl, H. W., & Huisman, J. (2016). How rising CO2 and global warming may stimulate harmful cyanobacterial blooms. Harmful Algae, 54, 145–159. 10.1016/j.hal.2015.12.006

85 Wanner, B. L. (1993). Gene regulation by phosphate in enteric bacteria. Journal of Cellular Biochemistry, 51(1), 47–54. 10.1002/jcb.240510110

86 Watkins, S. C., Smith, J. R., Hayes, P. K., & Watts, J. E. M. (2014). Characterisation of Host Growth after Infection with a Broad-Range Freshwater Cyanopodophage. PLOS ONE, 9(1), e87339. 10.1371/journal.pone.0087339

87 Weitz, J. S., & Wilhelm, S. W. (2012). Ocean viruses and their effects on microbial communities and biogeochemical cycles. F1000 Biology Reports, 4, 17. 10.3410/B4-17

88 Wilson, A. E., Wilson, W. A., & Hay, M. E. (2006). Intraspecific variation in growth and morphology of the bloom-forming cyanobacterium *Microcystis aeruginosa*. Applied and Environmental Microbiology, 72(11), 7386–7389. 10.1128/AEM.00834-06

89 Winter, S. E., Thiennimitr, P., Winter, M. G., Butler, B. P., Huseby, D. L., Crawford, R. W., Russell, J. M., Bevins, C. L., Adams, L. G., Tsolis, R. M., Roth, J. R., & Bäumler, A. J. (2010). Gut inflammation provides a respiratory electron acceptor for Salmonella. Nature, 467(7314), Article 7314. 10.1038/nature09415

90 Wucher, B. R., Winans, J. B., Elsayed, M., Kadouri, D. E., & Nadell, C. D. (2023). Breakdown of clonal cooperative architecture in multispecies biofilms and the spatial ecology of predation. Proceedings of the National Academy of Sciences, 120(6), e2212650120. 10.1073/pnas.2212650120

91 Xiao, M., Willis, A., & Burford, M. A. (2017). Differences in cyanobacterial strain responses to light and temperature reflect species plasticity. Harmful Algae, 62, 84–93. 10.1016/j.hal.2016.12.008

92) Yancey, C. E., Smith, D. J., Den Uyl, P. A., Mohamed, O. G., Yu, F., Ruberg, S. A., Chaffin, J. D., Goodwin, K. D., Tripathi, A., Sherman, D. H., & Dick, G. J. (2022). Metagenomic and Metatranscriptomic Insights into Population Diversity of *Microcystis* Blooms: Spatial and Temporal Dynamics of mcy Genotypes, Including a Partial Operon That Can Be Abundant and Expressed. Applied and Environmental Microbiology, 88(9), e0246421. 10.1128/aem.02464-21

93 Yancey, C. E., Kiledal, E. A., Chaganti, S. R., Denef, V. J., Errera, R. M., Evans, J. T., Hart, L. N., Isailovic, D., James, W. S., Kharbush, J. J., Kimbrel, J. A., Li, W., Mayali, X., Nitschky, H., Polik, C. A., Powers, M. A., Premathilaka, S. H., Rappuhn, N. A., Reitz, L. A., … Dick, G. J. (2023). The Western Lake Erie culture collection: A promising resource for evaluating the physiological and genetic diversity of *Microcystis* and its associated microbiome. Harmful Algae, 126, 102440. 10.1016/j.hal.2023.102440

94 Yoshida, M., Yoshida, T., Kashima, A., Takashima, Y., Hosoda, N., Nagasaki, K., & Hiroishi, S. (2008). Ecological dynamics of the toxic bloom-forming cyanobacterium *Microcystis aeruginosa* and its cyanophages in freshwater. Applied and Environmental Microbiology, 74(10), 3269–3273. 10.1128/AEM.02240-07

95 Yoshida, T., Takashima, Y., Tomaru, Y., Shirai, Y., Takao, Y., Hiroishi, S., & Nagasaki, K. (2006). Isolation and Characterization of a Cyanophage Infecting the Toxic Cyanobacterium *Microcystis aeruginosa*. Applied and Environmental Microbiology, 72(2), 1239–1247. 10.1128/AEM.72.2.1239-1247.2006

96 Yoshida-Takashima, Y., Yoshida, M., Ogata, H., Nagasaki, K., Hiroishi, S., & Yoshida, T. (2012). Cyanophage infection in the bloom-forming cyanobacteria *Microcystis aeruginosa* in surface freshwater. Microbes and Environments, 27(4), 350–355. 10.1264/JSME2.ME12037

97) Zhu, L., Zancarini, A., Louati, I., De Cesare, S., Duval, C., Tambosco, K., Bernard, C., Debroas, D., Song, L., Leloup, J., & Humbert, J.-F. (2016). Bacterial Communities Associated with Four Cyanobacterial Genera Display Structural and Functional Differences: Evidence from an Experimental Approach. Frontiers in Microbiology, 7. https://www.frontiersin.org/articles/10.3389/fmicb.2016.01662

98 Zimmerman, A. E., Howard-Varona, C., Needham, D. M., John, S. G., Worden, A. Z., Sullivan, M. B., Waldbauer, J. R., & Coleman, M. L. (2020). Metabolic and biogeochemical consequences of viral infection in aquatic ecosystems. Nature Reviews Microbiology, 18(1), Article 1. 10.1038/s41579-019-0270-x

99 Zinser, E. (2018). The microbial contribution to reactive oxygen species dynamics in marine ecosystems. Environmental Microbiology, 10(4). https://doi-org.proxy.lib.umich.edu/10.1111/1758-2229.12626

